# Genome-Wide Association Study and Pathway-Level Analysis of Kernel Color in Maize

**DOI:** 10.1101/535872

**Authors:** Brenda F. Owens, Deepu Mathew, Christine H. Diepenbrock, Tyler Tiede, Di Wu, Maria Mateos-Hernandez, Michael A. Gore, Torbert Rocheford

## Abstract

Rapid development and adoption of biofortified, provitamin A-dense orange maize (*Zea mays* L.) varieties could be facilitated by a greater understanding of the natural variation underlying kernel color, including as relates to carotenoid biosynthesis and retention in maize grain. Greater abundance of carotenoids in maize kernels is generally accompanied by deeper orange color, useful for distinguishing provitamin A-dense varieties to consumers. While kernel color can be scored and selected with high-throughput, low-cost phenotypic methods within breeding selection programs, it remains to be well established as to what would be the logical genetic loci to target for selection for kernel color. We conducted a genome-wide association study of maize kernel color, as determined by colorimetry, in 1,651 yellow and orange inbreds from the Ames maize inbred panel. Associations were found with *y1*, encoding the first committed step in carotenoid biosynthesis, and with *dxs2*, which encodes the enzyme responsible for the first committed step in the biosynthesis of the isoprenoid precursors of carotenoids. These genes logically could contribute to overall carotenoid abundance and thus kernel color. The *lcyE* and *zep1* genes, which can affect carotenoid composition, were also found to be associated with colorimeter values. A pathway-level analysis, focused on genes with *a priori* evidence of involvement in carotenoid biosynthesis and retention, revealed associations for *dxs3* and *dmes1*, involved in isoprenoid biosynthesis; *ps1* and *vp5*, within the core carotenoid pathway; and *vp14*, involved in cleavage of carotenoids. Collectively, these identified genes appear relevant to the accumulation of kernel color.

## INTRODUCTION

Malnutrition, or hidden hunger, remains a serious issue, even as increased agricultural productivity has helped to provide more energy and calories on a global scale (Welch and Graham 1999). As much as half of the world’s population may be deficient in one or more micronutrients, with 125–130 million pre-school children and 7 million pregnant women suffering from vitamin A deficiency (VAD) (Stevens *et al*. 2015). Biofortification, the improvement of crop nutritional quality through breeding and/or agronomics, has been proposed as a sustainable tool to help with addressing micronutrient malnutrition (Bouis and Welch 2010), and has found to be cost-effective (Meenakshi *et al*. 2012; Bouis and Hunt 1999; Qaim *et al*. 2007). Improvement of provitamin A carotenoid levels is generally a promising target, given that naturally occurring yellow and orange-pigmented accessions have been identified for many white-pigmented, starchy staple foods such as maize, cassava, banana, and sweet potato (Amorim *et al*. 2009; Carvalho *et al*. 2016; Takahata *et al*. 1993).

For biofortification to be effective, micronutrient densities must reach levels that impact human health, and the varieties and final food products must be acceptable to growers and consumers. Through decades of technical and broader contextual work, the international breeding organizations of CIMMYT, IITA and HarvestPlus, and partners have achieved the successful development of provitamin A-dense maize varieties, nearing target nutrient levels, which also have local and regional adaptation and relevance (Pixley *et al*. 2013, Menkir *et al*. 2017). Specifically, there has been a need to develop maize with distinctly orange kernel color for enhanced product recognition and enhanced consumer acceptance, including in certain sub-Saharan African nations where white maize is preferred but outreach and educational initiatives have successfully linked enhanced nutritional properties to the novel orange color (Meenakshi *et al*. 2012; Muzhingi *et al*. 2008; reviewed in Simpungwe *et al*. 2017). For the consistent and facilitated development of biofortified, provitamin A-dense maize varieties that meet target nutrient levels and also have strongly orange endosperm, it is important to identify and dissect the genetic loci underlying kernel color, including as relates to carotenoid content and composition. Relatedly, genetic loci showing consistent associations with darker orange color could in turn be targets for marker-assisted selection (MAS), in parallel with selection for provitamin A levels (Harjes *et al*. 2008; Yan *et al*. 2010; Menkir *et al*. 2012) and improved or maintained agronomic performance (Bouis and Welch 2010, Pixley *et al*. 2013, Menkir *et al*. 2017).

Carotenoids, including the provitamin A compounds α-carotene, β-carotene, and β-cryptoxanthin, are members of a large group of isoprenoid compounds synthesized in plants. Deoxy-xylulose 5-phosphate (DOXP) is formed by deoxy-xylulose 5-phosphate synthase (DXS) in the first step of the isoprenoid pathway. Seven more reactions are needed for the formation of the immediate carotenoid precursor, geranylgeranyl pyrophosphate (GGPP) from isopentenyl pyrophosphate (IPP) (Figure 1) (Hirschberg 2001; Rodríguez-Concepción and Boronat 2002; Hunter 2007; Rodríguez-Concepción *et al*. 2013; Vranová *et al*. 2013). The first committed step in carotenoid biosynthesis involves the formation of phytoene from two molecules of GGPP by phytoene synthase (PSY) (Buckner *et al*. 1996). Four more steps result in the biosynthesis of lycopene, after which there resides a key branch point in the pathway. For the biosynthesis of α-branch carotenoids, lycopene can be cyclized by lycopene β-cyclase (LCYB) at one end and by lycopene ε-cyclase (LCYE) at the other end to form α-carotene; from there, hydroxylation of the β-ring produces zeinoxanthin, and subsequent hydroxylation of the ε-ring produces lutein. Alternatively, for the biosynthesis of β-branch carotenoids, lycopene can be cyclized by LCYB at both ends to form β-carotene; from there, hydroxylation of one β-ring produces β-cryptoxanthin, and subsequent hydroxylation of the other β-ring produces zeaxanthin. Zeaxanthin can be further epoxidated to antheraxanthin and violaxanthin (Figure 2) (Hirschberg 2001; DellaPenna and Pogson 2006). A number of apocarotenoid metabolites are additionally formed from the oxidative cleavage of carotenoids by carotenoid cleavage dioxygenases (CCDs) and 9-cis-epoxycarotenoid dioxygenases (NCEDs), including strigolactones, abscisic acid (ABA), and various aromatic volatile compounds (Tan *et al*. 1997; Schwartz *et al*. 1997; Schwartz *et al*. 2001; Matusova *et al*. 2005; Sun *et al*. 2008; Vogel *et al*. 2008; Messing *et al*. 2010; Vallabhaneni *et al*. 2010; reviewed by Auldridge *et al*. 2006).

**Figure 1.**
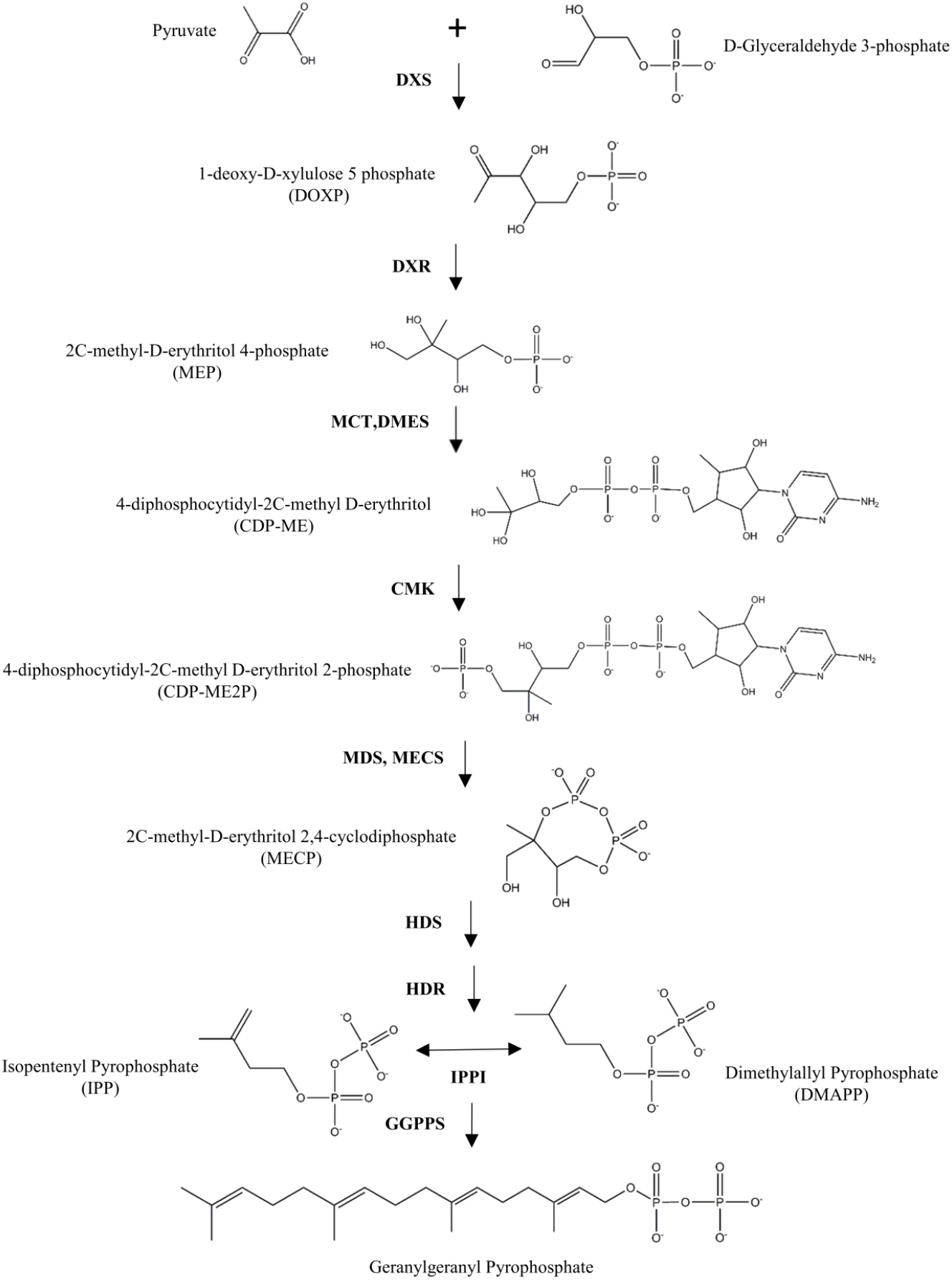
The methylerythritol 4-phosphate (MEP) biosynthetic pathway in plants. Compound names and abbreviations are as listed in the figure. Enzyme names and abbreviations: DOXP synthase (DXS), DOXP reductoisomerase (DXR), MEP cytidyltransferase (MCT), CDP-ME synthase (DMES), DP-ME kinase (CMK), MECP synthase (MDS, MECS), 4-hydroxy-3-methylbut-2-enyl-diphosphate [HMBBP] synthase (HDS), HMBBP reductase (HDR), isopentenyl pyrophosphate isomerase (IPPI), geranylgeranyl pyrophosphate synthase (GGPS).

**Figure 2.**
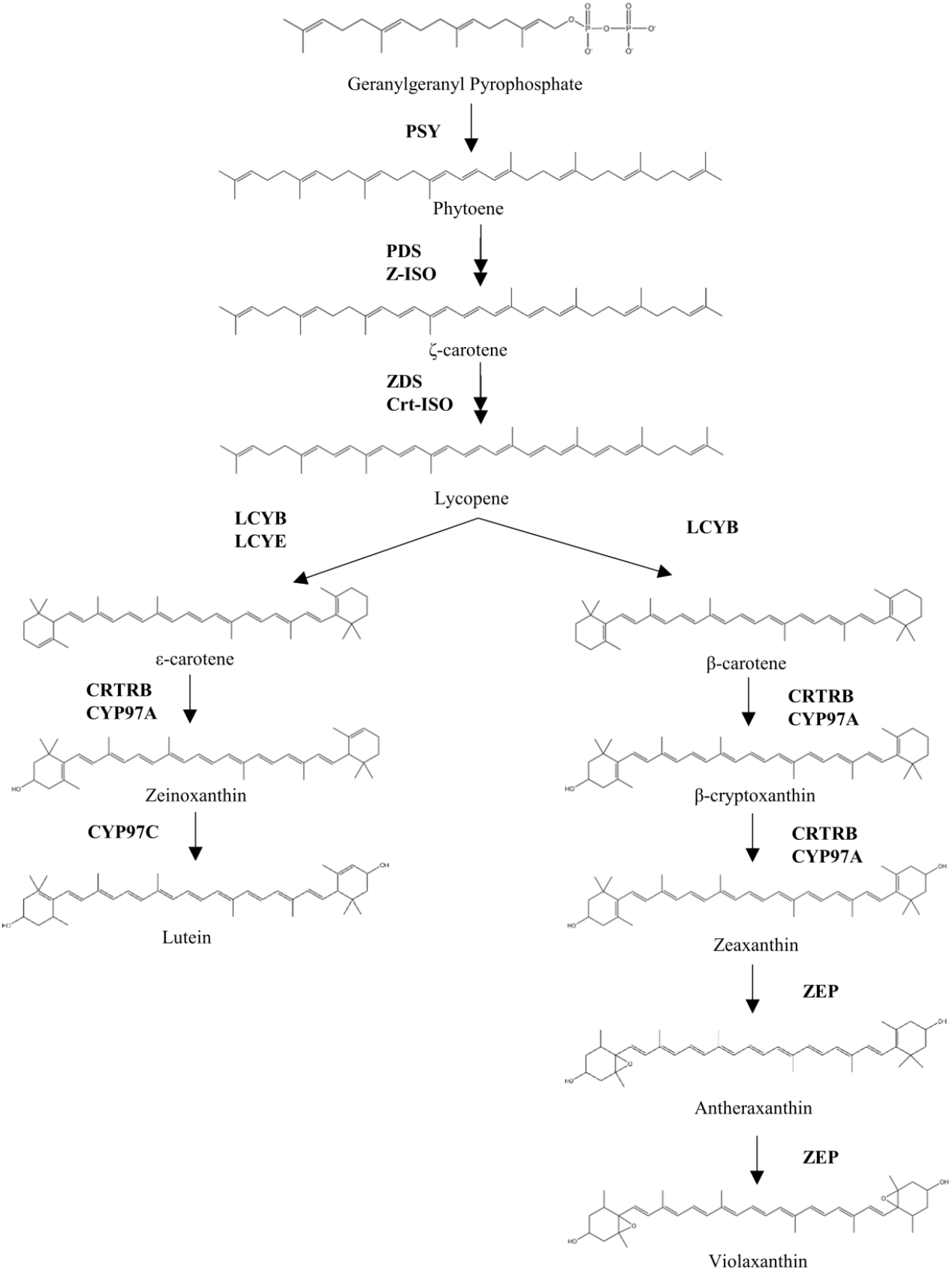
The carotenoid biosynthetic pathway in plants. Enzyme names and abbreviations: phytoene synthase (PSY), phytoene desaturase (PDS), *ζ*-carotene isomerase (ZDS), *ζ*-carotene desaturase (ZDS), carotenoid isomerase (Crt-ISO), lycopene β-cyclase (LCYB), lycopene ε-cyclase (LCYE), β-carotene hydroxylase (CRTRB), P450 carotenoid β-ring hydroxylase (CYP97A), P450 carotenoid epsilon-ring hydroxylase (CYP97C), zeaxanthin epoxidase (ZEP).

Many carotenoid compounds have yellow-to-red coloration dependent on functional groups and the length of their conjugated double bond systems (Khoo *et al*. 2011). Lutein and zeaxanthin, the two most abundant carotenoid compounds in maize grain (Owens *et al*. 2014), have been reported as light yellow and yellow-orange, respectively (Weber 1987; Meléndez-Martínez *et al*. 2007). Within the maize kernel, carotenoids predominantly accumulate in the vitreous portion of the endosperm (Weber 1987), though ABA which is derived from carotenoids plays a key role in the embryo in seed dormancy (McCarty 1995; Kermode 2005). The genes described in the isoprenoid and carotenoid biosynthetic pathways are logical *a priori* candidates for the genetic control of kernel color, given that their gene action could feasibly impact the hue and/or intensity of maize endosperm coloration.

Carotenoid composition, or relative abundance of individual carotenoid compounds, is typically quantified using high-performance liquid chromatography (HPLC). However, HPLC is cost- and labor-intensive and may not be amenable to the high-throughput measurements called for in certain stages of a breeding program (Diepenbrock and Gore 2015). For example, measurement methodologies that are still quantitative but less resource-intensive may have particular utility in the initial stages of breeding, in which large numbers of progeny are typically evaluated (Jaramillo *et al*. 2018, Ikeogu *et al*. 2017). However, it is important to understand the genetic loci underlying kernel color traits, particularly if colorimetry is to be used as a pre-screening tool in early-stage selections, so as not to select against favorable alleles at loci controlling provitamin A levels (or other compositional traits of importance to human health and nutrition).

Gradation in orange kernel color was previously visually scored on an ordinal scale, on bulks of kernels sampled from maize ears of 10 recombinant inbred line (RIL) families of the U.S. maize nested association mapping (NAM) population (McMullen *et al*. 2009). This study identified QTL for kernel color, which also mapped to regions containing carotenoid biosynthetic pathway genes (Chandler *et al*. 2013). Three QTL studies in other cereals identified intervals that were associated with colorimeter measurements of wheat endosperm, wheat flour, and sorghum endosperm, and that were in the vicinity of genes with putative involvement in carotenoid accumulation (Fernandez *et al*. 2008, Blanco *et al*. 2011, Zhao *et al*. 2013). These findings, combined with the rapid, inexpensive, and quantitative nature of colorimetric measurements, suggest that colorimetry may be a feasible method for quantification of maize kernel color in breeding programs, including for genetic analyses.

A colorimeter is an instrument that converts reflectance measurements into values that correspond to human perception of color. The CIELAB (*L*a*b\**) system is based on color-opponent theory, or color being perceived by the following pairs of opposites (Hunter and Harold 1987). The L axis represents a light to dark scale where positive values are lighter and negative values are darker. The “a” axis represents a greenness to redness scale where positive values are more red and negative values are more green. The “b” axis represents a yellowness to blueness scale where positive values are more yellow and negative values are more blue. Chroma is calculated from “a” and “b” values (Berger-Schunn 1994). Chroma represents the saturation or vividness of color, and hue represents the basic perceived color (whether the color would be called green or orange, for example) (Darrigues *et al*. 2008). Thus, hue and Chroma convert the *a\** and b* values to scores that represent a place in the color space to which humans have assigned a color name. Colorimeter values offer certain advantages over visual scoring given that they are quantitative, providing a more continuous scale of measurement; objective, allowing values to be compared across breeding populations over time; and representative of multiple components of kernel color.

Colorimetric methods were used in this study to genetically dissect the kernel color of 1,651 inbred lines from the Ames maize inbred panel (Romay *et al*. 2013). This study was conducted to 1) investigate the regions of the maize genome influencing kernel color using a genome-wide association study (GWAS), and 2) determine whether pathway-level analysis reveals additional associations with carotenoid-related genes.

## MATERIALS AND METHODS

### Experimental Design and Phenotypic Data

We grew a 2,448 experimental inbred line subset of a population consisting of 2,815 maize inbred lines maintained by the National Plant Germplasm System (Romay *et al*. 2013), hereafter referred to as the Ames maize inbred panel. Seed was provided by the North Central Regional Plant Introduction Station (NCRPIS) in Ames, IA, and grown as a single replicate at the Purdue Agronomy Center for Research and Education (ACRE) in West Lafayette, IN, in 2012 and 2013. The inbred lines were grouped into six sets based on maturity (i.e., flowering time) to facilitate pollination, harvesting and phenotyping efforts. Each set was arranged in a 20 x 24 incomplete block design. Each block within each set was augmented with an experiment-wide check line (B73) plot in a random position, and six other check lines of varying maturities based on flowering time (P39, Mo17, B97, NC358, Mo18W, CML247 in 2012 and PHJ40, Mo17, PHG35, PHG39, CML247, DK311H6 in 2013) were included twice per block in random positions. An experimental unit consisted of a one-row plot, 3.81 m in length containing approximately 15 plants. Plots had a spacing between rows of 0.762 m. Efforts were made to hand-pollinate up to six plants per plot. Self-pollinated ears were hand-harvested and dried for 72 h with forced hot air.

Inbred lines that were sweet corn or popcorn, or with white, red or blue endosperm color were removed from the data set because the kernels have characteristics that interfere with comparison of color measurements. Red and blue lines have pericarp color due to anthocyanins that are unrelated to carotenoid content, and white lines have very little carotenoid content. Popcorn and sweet corn have different kernel shapes than dent corn that may alter reflectance. This removal process resulted in 1,769 yellow and orange inbreds from the Ames panel that were analyzed by colorimetry.

To quantify kernel color, a Konica Minolta CR-400 Chroma Meter was used. This instrument is also called a colorimeter by the manufacturer and is described to perform colorimetry (https://sensing.konicaminolta.asia/product/chroma-meter-cr-400/). We will use the term colorimeter and colorimetry henceforth. The color values *L*, a*, b\**, and hue (*h*) were measured. Chroma (*C\**) values were not provided by the colorimeter, thus this value was calculated according to the formula Chroma = (*a*^2^ + b*^2^*)^1/2^. These measurements and calculated values correspond to the CIELAB *L*a*b\** system and the *L*C*h* system mathematically derived from it. Colorimeter settings used the standard illuminant D65 and an observer angle of 2° during the measurements. Three well-filled maize ears per plot were measured, with five random positions on each ear used for colorimeter recordings. The colorimeter was calibrated relative to a white reference before beginning measurements, and again every fifteen minutes while measurements were conducted.

### Phenotypic Data Analysis

To identify and remove significant outliers, a mixed linear model was fitted for each kernel colorimeter trait in ASReml-R version 3.0 (Gilmour *et al*. 2009). The full model fitted to the data was as follows:

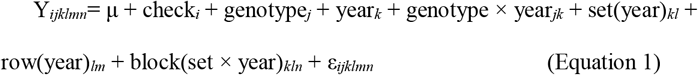

in which *Y_ijklmn_* is an individual phenotypic observation; μ is the grand mean; checki is the effect for the *i*th check; genotype_*j*_ is the effect of the *j*th experimental genotype (non-check line); year_*k*_ is the effect of the *k*th year; genotype × year_*jk*_ is the effect of the interaction between the *j*th genotype and *k*th year; set(year)_*kl*_ is the effect of the *l*th set within the *k*th year; row(year)_*lm*_ is the effect of the *m*th row within the *l*th year; block(set × year)_*kln*_ is the effect of the *n*th block within the *l*th set within the *k*th year; and ε_*ijklmn*_ is the residual error effect assumed to be independently and identically distributed according to a normal distribution with mean zero and variance *σ*_ε_^2^; that is, ~iid *N*(0, *σ*_ε_^2^). Degrees of freedom were calculated using the Kenward-Roger approximation (Kenward and Roger 1997). Except for the grand mean and check term, all other terms were fitted as random effects according to ~iid *N*(0, *σ*^2^). Studentized deleted residuals (Neter *et al*. 1996) generated from these mixed linear models were examined to remove significant outlier observations based on the Bonferroni correction at the α = 0.05 level of significance. Once significant outliers were removed, plot-level averages were calculated for each trait.

For each trait, the 2012 and 2013 plot-level averages were used in an iterative mixed linear model fitting procedure with the full model (Equation 1) in ASReml-R version 3.0 (Gilmour *et al*. 2009). Likelihood ratio tests were conducted to remove all terms fitted as random effects from the model that were not significant at α = 0.05 (Littell *et al*. 2006) to produce a final, best fitted model for each trait. For each trait, the final model was used to generate a best linear unbiased predictor (BLUP) for each genotype (Table S1).

Variance component estimates from the full model were used for estimation of heritability on a line-mean basis (Hung *et al*. 2012; Holland *et al*. 2003), with standard errors of the heritability estimates calculated using the delta method (Holland *et al*. 2003). Pearson’s correlation coefficient (r) was used to assess the degree of association between the BLUP values for each pair of colorimeter traits at α = 0.05 using the function ‘cor’ in R version 3.5.1 (R Core Team, 2018).

Prior to conducting the GWAS, the Box-Cox power transformation (Box and Cox 1964) was used on the BLUP values for each trait to correct for unequal error variances and non-normality of error terms (Table S2). The Box-Cox procedure was performed using the MASS package version 7.3-50 in R, with the evaluated lambda values ranging from −2 to +2 in increments of 0.5 to select the optimal convenient lambda for each trait. A lambda value of ‘2’ (square transformation) was obtained for hue and *L\**, whereas a lambda value of ‘1’ (no transformation) was obtained for *a*, b\**, and *C\**.

### Genome-wide association study

A GWAS was conducted for each of the five traits using the single-nucleotide polymorphism (SNP) data set developed using the genotyping-by-sequencing (GBS) platform for the Ames panel (Romay *et al*. 2013). The GBS marker data set used in this study consisted of partially imputed SNP genotypic data with B73 AGPv4 coordinates (ZeaGBSv27_publicSamples_imputedV5_AGPv4-161010.h5, available at panzea.org). Additional quality filters were imposed to retain SNPs with a call rate greater than 70%, minor allele frequency (MAF) greater than 2%, and inbreeding coefficient greater than 80%, resulting in a final dataset of 268,006 high-quality SNPs. In addition, inbred lines with lower than a 40% call rate were excluded.

For each kernel colorimeter trait, the GWAS was conducted using a mixed linear model that included the population parameters previously determined (Zhang *et al*. 2010) to test for an association between genotypes of each of the 268,006 SNPs and BLUP values from the 1,651 experimental inbred lines having both genotypic and phenotypic data, including after the above-described quality control steps. The GWAS was conducted in the R package GAPIT version 2017.08.18 (Lipka *et al*. 2012). To control for population structure and familial relatedness, the fitted mixed linear models included principal components (PCs) (Price *et al*. 2006) and a kinship matrix based on VanRaden’s method 1 (VanRaden 2008) calculated with the full set of 268,006 partially imputed SNPs. Before performing the GWAS, the missing genotypes remaining for all SNP markers were conservatively imputed as a ‘middle’ (heterozygous) value. The Bayesian information criterion (Schwarz 1978) was used to determine the optimal number of PCs to include as covariates in the mixed linear model for each trait. The amount of phenotypic variation accounted for the model was estimated with a likelihood-ratio-based *R*^2^ statistic (*R*^2^_LR_) (Sun *et al*. 2010). The Benjamini–Hochberg procedure (Benjamini and Hochberg 1995) was used to control the false discovery rate (FDR) at 5%.

### Pathway-level analysis

A set of 58 genes related to the biosynthesis and retention of carotenoids in maize was determined based on homology with known genes in *Arabidopsis thaliana*, and was previously used for a pathway-level analysis of carotenoid HPLC measurements in a small (*n* = 201) maize association panel (Owens *et al*. 2014). These same 58 genes, with the addition of *ζ*-carotene isomerase (*z-iso*) and homogentisate solanesyl transferase (w3), are referred to as pathway genes or *a priori* candidate genes in this study. Pathway-level analysis was used to reduce the number of association tests conducted, thus using *a priori* knowledge of the pathway to reduce the magnitude of the correction used to control the FDR at 5% (Califano *et al*. 2012; Owens *et al*. 2014). The set of 2,339 SNPs within ± 50 kb of the coding regions of the 60 *a priori* candidate genes was used in pathway-level analysis. The interval of ± 50 kb was a conservative estimate based on a previous finding in the Ames maize inbred panel of rapid decay of mean linkage disequilibrium in genic regions, reaching an average *r*^2^ = 0.2 within 1 kb, with large variance due to population structure, among other factors (Romay *et al*. 2013).

### Data availability

Phenotypes are provided in Tables S1 and S2 in the form of untransformed and transformed BLUPs. The GBS sequencing data are available at NCBI SRA (study accession number SRP021921). The SNP marker data are available at panzea.org, and accession names are listed in Tables S1 and S2.

## RESULTS

All of the colorimeter traits were highly heritable, with line-mean heritabilities ranging from 0.75 to 0.89 (Table 1). Hue values were positively correlated with *L\** (*r* = 0.75) and negatively correlated with *a\** (*r* = −0.94). Chroma and *b\** values were strongly positively correlated (*r* = 0.99) (Table 2). This correlation is likely due to *b\** values contributing most to Chroma (intensity of color), given the larger magnitude of *b\** relative to *a\** and the equal weighting of these two traits in the calculation of Chroma, whereas *a\** values corresponded more to hue (perceived color) in this data set.

**Table 1.**
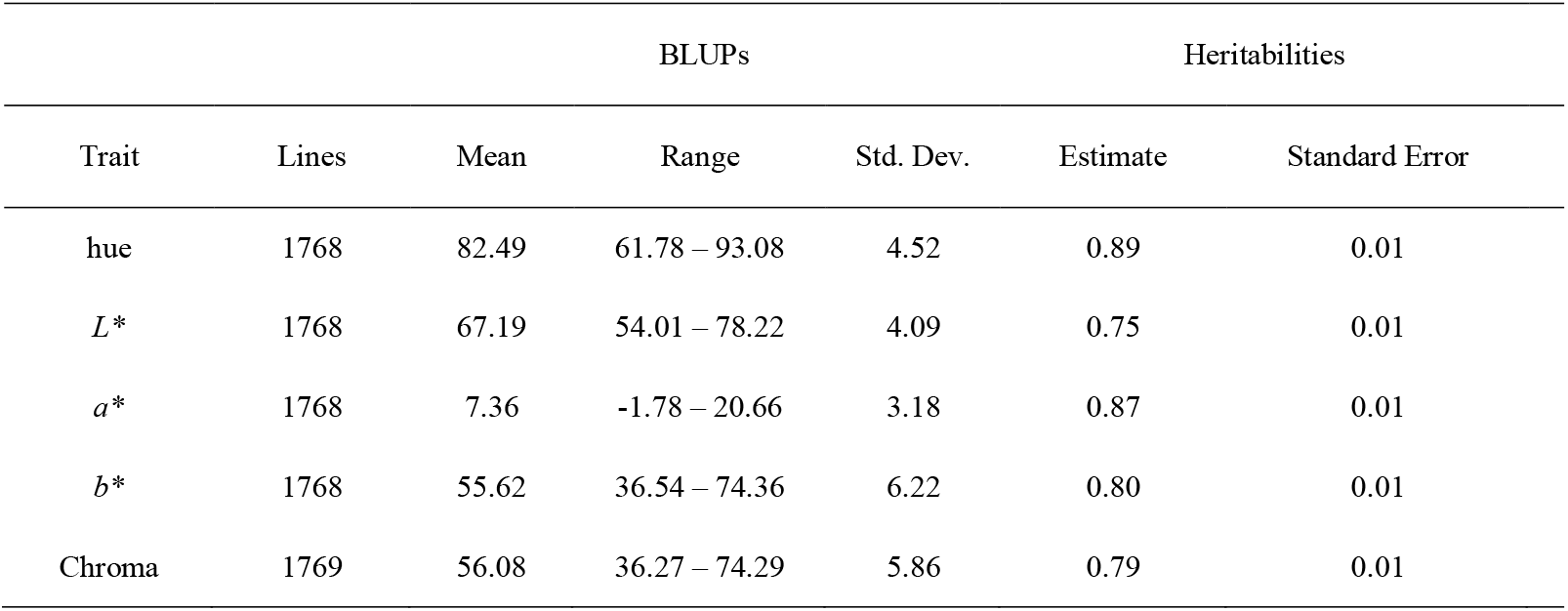
Means, ranges, and standard deviations (Std. Dev.) of untransformed BLUP values for five kernel colorimeter traits evaluated in the Ames maize inbred panel and estimated heritability on a line-mean basis across two years.

| Trait | Lines | BLUPs |  |  | Heritabilities |  |
| --- | --- | --- | --- | --- | --- | --- |
|  |  | Mean | Range | Std. Dev. | Estimate | Standard Error |
| hue | 1768 | 82.49 | 61.78 – 93.08 | 4.52 | 0.89 | 0.01 |
| *L\** | 1768 | 67.19 | 54.01 – 78.22 | 4.09 | 0.75 | 0.01 |
| *a\** | 1768 | 7.36 | -1.78 – 20.66 | 3.18 | 0.87 | 0.01 |
| *b\** | 1768 | 55.62 | 36.54 – 74.36 | 6.22 | 0.80 | 0.01 |
| Chroma | 1769 | 56.08 | 36.27 – 74.29 | 5.86 | 0.79 | 0.01 |

**Table 2.** Pearson’s correlation coefficients of untransformed BLUP values for five kernel colorimeter traits evaluated in the Ames maize inbred panel.

| | hue | $L^*$ | $a^*$ | $b^*$ | Chroma |
| --- | --- | --- | --- | --- | --- |
| hue | 1.00 | 0.75 | -0.94 | 0.54 | 0.44 |
| $L^*$ | | 1.00 | -0.68 | 0.58 | 0.52 |
| $a^*$ | | | 1.00 | -0.28 | -0.18 |
| $b^*$ | | | | 1.00 | 0.99 |
| Chroma |  |  |  |  | 1.00 |

All correlations were significant at $\alpha = 0.01$ .

A total of 27 unique SNPs were identified in GWAS for the five kernel colorimeter traits at an FDR-adjusted *P*-value of 5% (Table S3). Associations were detected for two genes involved in the provision of substrate for carotenoid biosynthesis. A single SNP was detected within (i.e., in the coding region of) a gene encoding 1-deoxy-D-xylulose 5-phosphate synthase (*dxs2*), the first and committed step in the isoprenoid biosynthetic pathway, with significant associations for *a\** and hue (Table 3). Two SNPs significantly associated with *a\** were detected within a gene encoding phytoene synthase (*y1*), the first and committed step in the biosynthesis of carotenoids.

**Table 3.** Carotenoid-related genes identified through genome-wide association study of five kernel colorimeter traits in the Ames maize inbred panel, and the most significant SNP for each trait-by-gene combination.

| Gene ID | Gene | Trait | SNP ID | Chr | Position of SNP | P-value | FDR-adjusted P-value | MAF | $R^2_{LR}$ | $R^2_{LR-SNP}$ |
| --- | --- | --- | --- | --- | --- | --- | --- | --- | --- | --- |
| Zm00001d003512 | *zep1* | *a* * | S2_44473748 | 2 | 46,329,893 | 7.30E-07 | 2.44E-02 | 0.141 | 0.414 | 0.423 |
| Zm00001d036345 | *y1* | *a* * | S6_82020346 | 6 | 85,064,521 | 2.65E-07 | 1.58E-02 | 0.061 | 0.414 | 0.423 |
| Zm00001d019060 | *dxs2* | *a* * | S7_14078791 | 7 | 14,495,640 | 3.25E-10 | 8.72E-05 | 0.050 | 0.414 | 0.428 |
| Zm00001d019060 | *dxs2* | hue | S7_14078791 | 7 | 14,495,640 | 4.74E-08 | 6.35E-03 | 0.050 | 0.493 | 0.502 |
| Zm00001d011210 | *lcyE* | hue | S8_138888278 | 8 | 143,026,247 | 2.52E-07 | 2.25E-02 | 0.416 | 0.493 | 0.501 |

**Gene ID:** Gene designation and position of SNPs from B73 RefGen\_v4 (www.maizegdb.org; Andorf et al. 2016); **Gene:** Annotated gene containing SNP, except *zep1*, which is 25 kb upstream of the SNP; **Position of SNP:** Genomic position (bp) of the SNP from B73 Refgen\_v4; **FDR-adjusted P-value:** False discovery rate adjusted *P*-value; **MAF:** Minor-allele frequency; **$R^2_{LR}$ :** $R^2$ likelihood ratio value of model without SNP; **$R^2_{LR-SNP}$ :** $R^2$ likelihood ratio value of model with SNP.

Two genes in the core carotenoid pathway were also identified. Two significant SNP associations were detected for hue within the gene encoding lycopene ε-cyclase (*lcyE*), which affects the partitioning of substrate into the α- and β- branches of the carotenoid pathway. A significant SNP associated with *a\** was located near the gene encoding zeaxanthin epoxidase (*zep1*), approximately 25 kb downstream of the gene. Zeaxanthin epoxidase converts zeaxanthin to antheraxanthin and subsequently violaxanthin, all within the β-branch of the pathway.

Twenty-one SNPs having significant associations with one or more traits did not have an *a priori* candidate gene within the ± 50 kb search space. These search spaces were subsequently examined, in case they contained other genes having plausible biological involvement with kernel color. Briefly, three significant SNPs for *a\** were proximal to GRMZM2G063663 (chr. 1). The product of this gene model was found to have 96% identity at the protein level with cytochrome P450 14 (CYP14, encoded by *lut1*, GRMZM2G143202). Three other significant SNPs for *a\** were proximal to a gene that encodes isopentenyl transferase (*ipt10*, GRMZM2G102915, chr. 6) and is expressed in the endosperm of B73 (Andorf et al. 2016). IPT transfers the five-carbon isoprenoid moiety from DMAPP, an isomer of IPP (Figure 1), to a certain position on tRNAs. Finally, one significant SNP for *b\** and two significant SNPs for chroma were proximal to a gene encoding enolase (*enolase1, eno1*, GRMZM2G064302, chr. 9), the penultimate enzyme in glycolysis. This gene was highly expressed in endosperm of B73 (Andorf et al. 2016).

We conducted a pathway-level analysis in which only SNPs within ± 50 kb of an *a priori* gene for carotenoid biosynthesis and/or retention were tested. This analysis revealed additional associations for colorimeter traits with all four of the carotenoid genes identified in GWAS: two SNPs in the coding region of *dxs2*, four SNPs in the coding region of *y1*, nine SNPs in the coding region of *lcyE*, and three SNPs proximal to *zep1* (Table 4, Table S4).

**Table 4.** Most significant SNP for each trait-by-gene combination within 50 kb of carotenoid-related genes identified through pathway-level association analyses of five kernel colorimeter traits in the Ames maize inbred panel.

| Gene ID | Gene | Trait | SNP ID | Chr | Position of SNP | P-value | FDR-adjusted P-value | MAF | $R^2_{LR}$ | $R^2_{LR-SNP}$ |
| --- | --- | --- | --- | --- | --- | --- | --- | --- | --- | --- |
| Zm00001d027936 | *vp5* | *a* * | S1_17625344 | 1 | 17,930,201 | 6.16E-05 | 1.02E-02 | 0.185 | 0.414 | 0.420 |
| Zm00001d033222 | *vp14* | *L* * | S1_250915992 | 1 | 255,044,348 | 1.59E-05 | 3.68E-02 | 0.161 | 0.336 | 0.344 |
| Zm00001d003513 | *zep1* | *a* * | S2_44443991 | 2 | 46,299,837 | 9.89E-06 | 3.81E-03 | 0.401 | 0.414 | 0.421 |
| Zm00001d003513 | *zep1* | hue | S2_44443991 | 2 | 46,299,837 | 1.47E-04 | 2.83E-02 | 0.401 | 0.493 | 0.498 |
| Zm00001d042584 | *dmes1* | hue | S3_170106723 | 3 | 172,731,046 | 3.47E-04 | 4.72E-02 | 0.180 | 0.493 | 0.497 |
| Zm00001d015651 | *ps1* | *a* * | S5_100735811 | 5 | 103,264,914 | 2.53E-04 | 3.45E-02 | 0.096 | 0.414 | 0.419 |
| Zm00001d036345 | *y1* | *b* * | S6_82018091 | 6 | 85,062,266 | 2.85E-05 | 3.30E-02 | 0.062 | 0.288 | 0.295 |
| Zm00001d036345 | *y1* | Chroma | S6_82018091 | 6 | 85,062,266 | 1.38E-05 | 2.96E-02 | 0.062 | 0.253 | 0.262 |
| Zm00001d036345 | *y1* | *a* * | S6_82019628 | 6 | 85,063,803 | 1.83E-06 | 1.06E-03 | 0.040 | 0.414 | 0.422 |
| Zm00001d036345 | *y1* | hue | S6_82020346 | 6 | 85,064,521 | 3.12E-04 | 4.56E-02 | 0.061 | 0.493 | 0.497 |
| Zm00001d019060 | *dxs2* | *a* * | S7_14078791 | 7 | 14,495,640 | 3.25E-10 | 7.53E-07 | 0.050 | 0.414 | 0.428 |
| Zm00001d019060 | *dxs2* | hue | S7_14078791 | 7 | 14,495,640 | 4.74E-08 | 1.10E-04 | 0.050 | 0.493 | 0.502 |
| Zm00001d011210 | *lcyE* | *a* * | S8_138882509 | 8 | 143,020,478 | 1.87E-05 | 3.93E-03 | 0.397 | 0.414 | 0.420 |
| Zm00001d011210 | *lcyE* | hue | S8_138882509 | 8 | 143,020,478 | 8.60E-06 | 2.49E-03 | 0.397 | 0.493 | 0.499 |
| Zm00001d011210 | *lcyE* | *b* * | S8_138886754 | 8 | 143,024,723 | 6.91E-06 | 1.60E-02 | 0.366 | 0.288 | 0.297 |
| Zm00001d011210 | *lcyE* | Chroma | S8_138886754 | 8 | 143,024,723 | 2.56E-05 | 2.96E-02 | 0.366 | 0.253 | 0.262 |
| Zm00001d045383 | *dxs3* | Chroma | S9_20472920 | 9 | 20,252,034 | 6.15E-05 | 4.75E-02 | 0.179 | 0.253 | 0.261 |

**Gene ID:** Gene designation and position of SNPs from B73 RefGen\_v4 (www.maizegdb.org); **Gene:** Annotated gene containing SNP or within 50 kb of SNP; **Position of SNP:** Genomic position (bp) of the SNP from B73 Refgen\_v4; **FDR-adjusted P-value:**
False discovery rate adjusted $P$ -value; **MAF**: Minor-allele frequency; $R^2_{\text{LR}}$ : $R^2$ likelihood ratio value of model without SNP; $R^2_{\text{LR-SNP}}$ : $R^2$ likelihood ratio value of model with SNP.

Additional associations were identified through pathway analysis in regions proximal to a number of genes not identified in GWAS. An association was found for Chroma in the vicinity of another gene that encodes DXS (*dxs3*, chr. 9). Two SNPs were significant for hue in the vicinity of 4-diphosphocytidyl-2C-methyl-D-erythritol synthase (*dmes1*, chr. 3), another gene in the isoprenoid biosynthetic pathway. Within the core carotenoid pathway, two additional genes were identified for *a*:* lycopene β-cyclase (*lycB, ps1, vp7*, chr. 5) and phytoene desaturase (*vp5*, chr. 1). Finally, a gene related to carotenoid cleavage, encoding 9-cis-epoxycarotenoid dioxygenase (NCED) (*vp14*, chr. 1), was identified for *L\**.

## DISCUSSION

A colorimeter was used to quantify kernel color in a large, diverse maize inbred panel. Visual color scoring has shown effectiveness in biparental crosses, where only a few classes of kernel color are segregating (Chandler *et al*. 2013), but is not suitable or tractable for large diversity panels with continuous gradients of kernel color. The most significant association in this study was detected for a SNP in the coding region of *dxs2*—one of three genes in the maize genome encoding DXS, the first enzyme in the non-mevalonate plastidic isoprenoid pathway (Cordoba *et al*. 2011). Significant associations were also detected in the coding region of phytoene synthase (y1), a gene that controls the first committed step in carotenoid biosynthesis (Buckner *et al*. 1996; Cunningham and Gantt 1998; DellaPenna and Pogson 2006).

Although joint linkage analysis of visual color score data detected a QTL in the vicinity of *y1* (Chandler *et al*. 2013), neither *y1* nor *dxs2* were strong hits in a genome-wide association study of HPLC carotenoid data in 201 inbreds with yellow to orange kernel color from the Goodman-Buckler diversity panel (Owens *et al*. 2014). In the present study of kernel color in a large association panel of 1,651 inbreds, significant associations were detected in the coding regions of both of these genes. PSY has been considered to be the key enzyme limiting carotenoid accumulation in maize endosperm (Zhu *et al*. 2008). The identification of *dxs2* and *y1* in this study indicates that genetic variation at these loci is associated with kernel color, likely due to the role of these genes in substrate provision for the biosynthesis of pigmented carotenoids. These genes merit further examination given that *dxs2* and *y1* respectively encode the first and committed steps in the IPP (precursor) pathway and the core carotenoid pathway, and showed the most significant statistical associations in this study. In particular, investigation of the main effects and any interaction effects of these two genes in maize, as well as their expression dynamics through kernel development and upon the overexpression or knockdown of one or both genes, may provide further insight into the extent to which their association with kernel color (and potentially carotenoids) is separate versus coordinated.

Associations in the regions of *lcyE* and *zep1*—genes affecting flux within and through the core carotenoid pathway—were identified both in this study of kernel color and in the prior study of carotenoid HPLC values in the Goodman-Buckler panel (Owens *et al*. 2014). Notably, signals in the vicinity of three of the genes identified in our GWAS—*lcyE, zep1*, and *y1*—were also detected in a previous joint-linkage analysis of visual scores for gradation in orange kernel color in 10 families of the U.S. maize NAM population (Chandler *et al*. 2013). Carotenoid compounds in the α-vs. β-branches have different spectral properties that influence color, due to differing numbers of double bonds in their structures. Specifically, the β-branch compounds (β-carotene, β-cryptoxanthin, and zeaxanthin) have 11 conjugated double bonds and correspondingly have lower *a\** values and higher *b\** values than α-carotene and lutein, which have 10 conjugated double bonds (Meléndez-Martínez *et al*. 2007; Khoo *et al*. 2011). Thus, a shift in the relative concentrations of these compounds has the potential to affect color.

For *lcyE*, encoding a protein that acts at the key pathway branch point, associations were indeed seen in the Goodman-Buckler panel for two ratio traits (β-branch to α-branch carotenoids, and β-branch to α-branch xanthophylls) as well as lutein, zeaxanthin, total α-xanthophylls, and total β-xanthophylls. An allele of *lcyE* with reduced expression was found to result in the formation of fewer ε-rings and a reduction in α-branch compounds relative to β-branch compounds (Harjes *et al*. 2008). Similarly to *lcyE*, associations with *zep1*— encoding a protein that acts within the β-pathway branch—were seen in the Goodman-Buckler panel for the ratio trait of β-branch to α-branch xanthophylls, as well as zeaxanthin and total β-xanthophylls.

Taken together, the identification of *dxs2* and *y1* (genes involved in overall substrate provision) in the present study suggests that kernel color can be utilized to select for greater carotenoid abundance in general. However, the simultaneous identification of *lcyE* and *zep1* (genes involved in carotenoid composition) suggests that the relative abundance of individual carotenoid compounds is likely to also be affected when selecting on kernel color. Therefore, the levels of individual carotenoid compounds will need to be monitored when colorimetry is applied as an early selection tool for lines having favorable orange color, to ensure that the favorable genetic variants needed for the maintenance or improvement of provitamin A levels are also retained. For example, the concentrations of the more abundant provitamin A carotenoids in maize grain, β-carotene and β-cryptoxanthin, might be increased simultaneously with orange kernel color if substrate were to be modulated via *lcyE* to flow preferentially through the β-branch of the pathway. Alternatively or in addition, favorable alleles of the gene encoding β-carotene hydroxylase (*crtRB1*), which converts β-carotene to β-cryptoxanthin to zeaxanthin, could be selected that favor accumulation and retention of these provitamin A compounds while also producing sufficient zeaxanthin to obtain the vivid orange color.

While there are many cytochrome P450s in the maize genome, the high level of homology between the product of GRMZM2G063663 and CYP14, which acts within the α-branch of the carotenoid pathway, suggests that this gene is a candidate for further examination. Regarding isopentenyl transferase (IPT), its activity has been found in maize to affect the distribution of aleurone vs. starchy endosperm layers (Geisler-Lee and Gallie 2005). Certain aleurone-deficient mutants have been found to be deficient in carotenoids, and it has been suggested that there may be some functional connection between aleurone differentiation and carotenoid biosynthesis (reviewed in Gontarek and Becraft 2017). The finding of signals proximal to *ipt10* in this study for kernel color suggests a potential genetic target for the further investigation of that hypothesis. Finally, the product of enolase— phosphoenylpyruvate (PEP)—has many potential metabolic routes. Nevertheless, the action of enolase resides only two steps prior to that of DXS (which takes pyruvate as one of its substrates), and PEP is an important precursor for isoprenoid biosynthesis. An engineering strategy in *E. coli* that increased PEP concentrations was found to increase levels of lycopene, the carotenoid compound that sits at the pathway branch point (Zhang et al. 2013). While *enolase1* may have underlaid associations with kernel color in this GWAS, it may not be a viable breeding target given the relatively higher likelihood of complex and/or unfavorable pleiotropic effects within central metabolism.

The pathway-level analysis conducted in this study revealed a number of additional genes significantly associated with color. Notably, an association with *dxs3* suggests that this gene, in addition to *dxs2*, may play a role in the accumulation of carotenoids in the maize kernel. An association was found with *dmes2*, which encodes 4-diphosphocytidyl-2C-methyl-D-erythritol synthase, the third step in the MEP pathway. The gene encoding this enzyme in *A. thaliana*, present in a single copy and termed *MCT*, has been found along with certain other MEP pathway genes to have very low seed expression levels in certain developmental stages, in a manner that may be limiting to carotenoid biosynthesis (Meier *et al*. 2011). In this study, the associations with isoprenoid pathway genes are an indication that the genetic control of the provision of IPP, a precursor for biosynthesis of carotenoids and other isoprenoids, is relevant to kernel color.

Three genes underlying classical viviparous maize mutants were identified in this study: vp5, encoding PDS (Hable *et al*. 1998); *vp7*, encoding LCYB (Singh *et al*. 2003); and *vp14*, encoding NCED (Tan *et al*. 1997). These three genes were previously recognized as Class Two viviparous mutants, which in addition to vivipary (precocious germination) exhibit altered endosperm and seedling color due to effects on carotenoid and chlorophyll biosynthesis (Robertson 1955). These three mutants have also been found to be deficient in ABA (McCarty 1995; Schwartz *et al*. 1997). The action of PDS and LCYB takes place prior to and coincident with the pathway branch point, respectively. The two corresponding mutants are also deficient in carotenoids (McCarty 1995), which would tend to affect kernel color if the pigmented carotenoids are among those depleted. NCED acts within the β-pathway branch, cleaving 9-cis-xanthophylls to xanthoxin (Tan *et al*. 1997), which is then converted to ABA. The *vp14* mutant was found to have reduced levels of zeaxanthin compared to wild type, though levels of the immediate substrates of NCED were unaffected (Tan *et al*. 1997). Given the finding of an effect on zeaxanthin levels, and the general action of NCED in the portion of the pathway corresponding to pigmented β-branch carotenoids and their derivatives, the association of genetic variation at *vp14* with kernel color is not entirely surprising.

Another cleavage enzyme, CCD—encoded by one or more copies of *ccd1* within the *White Cap (Wc)* locus in maize (Tan *et al*. 2017)—was not detected as being associated with natural variation in this study. The *Wc* locus was created in some maize accessions by a macrotransposon insertion, with subsequent tandem duplications resulting in the amplification of *ccd1* copy number in a subset of those accessions, and has been found to impact endosperm color through the degradation of carotenoids by CCD. Notably, the *Wc* locus was likely identified in the previously conducted analysis of visual scores for gradation in orange kernel color in 10 U.S. maize NAM families. While the *ccd1* progenitor locus (*Ccd1r*) was not contained in the QTL support interval identified on chromosome 9 (149.54 to 151.48 Mb, AGP v2), the macrotransposon insertion that created *Wc* was subsequently characterized in Tan *et al*. (2017), and appears to have been included in the interval. This QTL putatively corresponding to *Wc* was only significant in two of the 10 NAM families analyzed (Chandler *et al*. 2013), suggesting the possibility of rare variation at the *Wc* locus which may have precluded its identification in the present study. Additionally, given the tandem duplications inherent to *Wc* in some accessions, potentially informative paralogous SNP markers in this region may have been excluded in the SNP filtering process in the present study. Alternatively, the localization of variation relating to CCD may have been dispersed at the genetic level among a varying number of *ccd1* copies within *Wc* (in addition to the *Ccd1r* progenitor locus itself). This dispersion could present particular difficulties for the detection of genetic signal in the presence of low SNP coverage and/or rare variation. Finally, given that only lines with yellow to orange endosperm were analyzed in this study, it could be that the variation in *ccd1* copy number was too constrained (with yellow-endosperm lines being on the lower end of the dynamic range in copy number; Tan *et al*. 2017) for a genetic association with loci encoding CCD to be present and/or identified in this panel.

Further studies are needed to determine whether natural variation at the loci identified in these analyses corresponds to differences in transcription levels, post-translational regulation, and/or enzyme activity. A GWAS using kernel color phenotypes and HPLC-based carotenoid values for the same set of materials may enable the identification of alleles that are favorable for kernel color as well as carotenoid composition and concentration. An increasing knowledge of the genetic mechanisms affecting kernel color, and the potential relationships between color values and carotenoid values, will be useful in coordinating breeding efforts to improve both sets of phenotypes. Establishing optimal ranges for each colorimeter value for use in a selection index could provide a useful and inexpensive breeding tool, particularly to screen for kernel color and total carotenoid levels in the early stages of breeding. Some of the evaluation, selection, and elimination could potentially be done while the ears are still on the plants, or in a harvest pile at the end of a nursery row. This would save labor and reduce handling of non-selected ears. Selection of favorable alleles of the loci detected in this study, particularly *y1* and *dxs2*, in conjunction with the previously established alleles of *lcyE* and *crtRB1*, provide a logical and promising strategy for the rapid development of provitamin A-dense maize lines that also produce a recognizable and desirable orange kernel color.

## ACKNOWLEDGEMENTS

This research was supported by the National Science Foundation (NSF) IOS-0922493 (TRR) and IOS-1546657 (MAG), Harvest Plus (TRR, MAG), Purdue Patterson Chair funds (TRR), Cornell University startup funds (MAG), and by a USDA National Needs Fellowship (CHD). We thank Jerry Chandler and Chris Hoagland for assistance with fieldwork and seed handling.

## REFERENCES

Amorim, E. P., A. D. Vilarinhos, K. O. Cohen, V. B. O. Amorim, J. A. dos Santos-Serejo et al., 2009 Genetic diversity of carotenoid-rich bananas evaluated by Diversity Arrays Technology (DArT). Genet. Mol. Biol. 32(1): 96–103.

Andorf, C. M., E. K. Cannon, J. L. Portwood, J. M. Gardiner, L. C. Harper et al., 2016 MaizeGDB update: new tools, data and interface for the maize model organism database. Nucleic Acids Res. 44: 1195–1201.

Auldridge, M. E., D. R. McCarty, and H. J. Klee, 2006 Plant carotenoid cleavage oxygenases and their apocarotenoid products. Curr. Opin. Plant Biol. 9(3): 315–321.

Benjamini, Y., and Y. Hochberg, 1995 Controlling the false discovery rate: a practical and powerful approach to multiple testing. J. Roy. Stat. Soc. B Met. 57: 289–300.

Berger-Schunn, A., 1994 Practical Color Measurement: A Primer for the Beginner, A Reminder for the Expert. Wiley, New York.

Blanco, A., P. Colasuonno, A. Gadaleta, G. Mangini, A. Schiavulli et al., 2011 Quantitative trait loci for yellow pigment concentration and individual carotenoid compounds in durum wheat. J. Cereal Sci. 54: 255–264.

Bouis, H., and J. Hunt, 1999 Linking food and nutrition security: past lessons and future opportunities. Asian Devel. Rev. 17(1,2): 168–213.

Bouis, H. E., and R. M. Welch, 2010 Biofortification—a sustainable agricultural strategy for reducing micronutrient malnutrition in the global south. Crop Sci. 50: S20–32.

Box, G. E. P., and D. R. Cox, 1964 An analysis of transformations. J. Roy. Stat. Soc. B Met. 26: 211–252.

Buckner, B., P. S. Miguel, D. Janick-Buckner, and J. L. Bennetzen, 1996 The y1 gene of maize codes for phytoene synthase. Genetics 143: 479–488.

Califano, A., A. J. Butte, S. Friend, T. Ideker, and E. Schadt, 2012 Leveraging models of cell regulation and GWAS data in integrative network-based association studies. Nat. Genet. 44: 841–847.

Carvalho, L. J. C. B., M. A. V. Agustini, J. V. Anderson, E. A. Vieira, C. R. B. de Souza et al., 2016 Natural variation in expression of genes associated with carotenoid biosynthesis and accumulation in cassava (Manihot esculenta Crantz) storage root. BMC Biol. 16(1): 133.

Chandler, K., A. E. Lipka, B. F. Owens, H. H. Li, E. S. Buckler et al., 2013 Genetic analysis of visually scored orange kernel color in maize. Crop Sci. 53: 189–200.

Cordoba, E., H. Porta, A. Arroyo, C. San Roman, L. Medina et al., 2011 Functional characterization of the three genes encoding 1-deoxy-D-xylulose 5-phosphate synthase in maize. J. Exp. Bot. 62: 2023–2038.

Cunningham, F. X., and E. Gantt, 1998 Genes and enzymes of carotenoid biosynthesis in plants. Annu. Rev. Plant Phys. and Plant Mol. Biol. 49: 557–583.

Darrigues, A., J. Hall, E. van der Knaap, D. M. Francis, N. Dujmovic, et al., 2008 Tomato Analyzer-Color Test: a new tool for efficient digital phenotyping. J. Amer. Soc. Hort. Sci. 133(4): 579–586.

DellaPenna, D., and B. J. Pogson, 2006 Vitamin synthesis in plants: tocopherols and carotenoids. Annu. Rev. Plant Biol. 57: 711–738.

Diepenbrock, C. H., and M. A. Gore, 2015 Closing the divide between human nutrition and plant breeding. Crop Sci. 55: 1437–1448.

Fernandez, M. G. S., M. T. Hamblin, L. Li, W. L. Rooney, M. R. Tuinstra et al., 2008 Quantitative trait loci analysis of endosperm color and carotenoid content in sorghum grain. Crop Sci. 48: 1732–1743.

Geisler-Lee, J., and D. R. Gallie, 2005 Aleurone cell identity is suppressed following connation in maize kernels. Plant Physiol. 139: 204–212.

Gilmour, A. R. G., B. B. Cullis, R. Thompson and D. Butler, 2009 ASReml user guide release 3.0. VSN International Ltd, Hemel Hempstead, UK.

Gontarek, B. C., and P. W. Becraft, 2017 Aleurone, pp. 68–80 in Maize Kernel Development, edited by B. A. Larkins. CABI, Boston.

Hable, W. E., K. K. Oishi, and K. S. Schumaker, 1998 Viviparous-5 encodes phytoene desaturase, an enzyme essential for abscisic acid (ABA) accumulation and seed development in maize. Mol. Gen. Genet. 257: 167–176.

Harjes, C. E., T. R. Rocheford, L. Bai, T. P. Brutnell, C. B. Kandianis et al., 2008 Natural genetic variation in lycopene epsilon cyclase tapped for maize biofortification. Science 319: 330–333.

Hirschberg, J., 2001 Carotenoid biosynthesis in flowering plants. Curr. Opin. Plant Biol. 4: 210–218.

Holland, J. B., W. E. Nyquist, and C. T. Cervantes-Martinez, 2003 Estimating and interpreting heritability for plant breeding: an update. Plant Breed. Rev. 22: 9–112.

Hung, H. Y., C. Browne, K. Guill, N. Coles, M. Eller et al., 2012 The relationship between parental genetic or phenotypic divergence and progeny variation in the maize nested association mapping population. Heredity 108: 490–499.

Hunter, R. S., and R. W. Harold, 1987 The Measurement of Appearance, 2nd ed. John Wiley & Sons, New York.

Hunter, W. N., 2007 The non-mevalonate pathway of isoprenoid precursor biosynthesis. J. Biol. Chem. 282: 21573–21577.

Ikeogu, U. N., F. Davrieux, D. Dufour, H. Ceballos, C. N. Egesi et al., 2017 Rapid analyses of dry matter content and carotenoids in fresh cassava roots using a portable visible and near infrared spectrometer (Vis/NIRS). PLoS ONE 12(12): e0188918.

Jaramillo, A. M., L. F. Londoño, J. C. Orozco, G. Patiño, J. Belalcazar et al., 2018 A comparison study of five different methods to measure carotenoids in biofortified yellow cassava (Manihot esculenta). PLoS ONE 13(12): e0209702.

Kenward, M. G., and J. H. Roger, 1997 Small sample inference for fixed effects from restricted maximum likelihood. Biometrics 53(3): 983–997.

Kermode, A. R., 2005 Role of abscisic acid in seed dormancy. J. Plant Growth Regul. 24: 319–344.

Khoo, H. E., K. N. Prasad, K. W. Kong, Y. Jiang and A. Ismail, 2011 Carotenoids and their isomers: color pigments in fruits and vegetables. Molecules 16: 1710–1738.

Lipka, A. E., F. Tian, Q. Wang, J. Peiffer, M. Li et al., 2012 GAPIT: genome association and prediction integrated tool. Bioinformatics 28: 2397–2399.

Littell, R. C., G. A. Milliken, W. W. Stroup, R. D. Wolfinger, and O. Schabenberger, 2006 SAS for Mixed Models. SAS Institute, Cary, N.C.

Matusova, R., K. Rani, F. W. A. Verstappen, M. C. R. Franssen, M. H. Beale, et al., 2005 The strigolactone germination stimulants of the plant-parasitic Striga and Orobanche spp. are derived from the carotenoid pathway. Plant Physiol. 139(2): 920–934.

McCarty, D. R., 1995 Genetic control and integration of maturation and germination pathways in seed development. Annu. Rev. Plant Physiol. Plant Mol. Biol. 46: 71–93.

McMullen, M. D., S. Kresovich, H. S. Villeda, P. Bradbury, H. H. Li et al., 2009 Genetic properties of the maize nested association mapping population. Science 325: 737–740.

Meenakshi, J. V., A. Banerji, V. Manyong, K. Tomlins, N. Mittal et al., 2012 Using a discrete choice experiment to elicit the demand for a nutritious food: willingness-to-pay for orange maize in rural Zambia. J. Health Econ. 31: 62–71.

Meier, S., O. Tzfadia, R. Vallabhaneni, C. Gehring, and E. T. Wurtzel, 2011 A transcriptional analysis of carotenoid, chlorophyll and plastidial isoprenoid biosynthesis genes during development and osmotic stress responses in Arabidopsis thaliana. BMC Syst. Biol. 5: 77.

Melendez-Martinez, A. J., G. Britton, I. M. Vicario, and F. J. Heredia, 2007 Relationship between the colour and the chemical structure of carotenoid pigments. Food Chem. 101: 1145–1150.

Menkir, A., B. Maziya-Dixon, W. Mengesha, T. Rocheford, and E. O. Alamu, 2017 Accruing genetic gain in pro-vitamin A enrichment from harnessing diverse maize germplasm. Euphytica 213: 105.

Menkir, A., K. Pixley, B. Maziya-Dixon, and M. Gedil, 2012 Recent advances in breeding maize for enhanced pro-vitamin A content, pp. 66–73 in Meeting the Challenges of Global Climate Change and Food Security through Innovative Maize Research: Proceedings of the Third National Maize Workshop of Ethiopia, edited by M. Worku, S. Twumasi-Afriyie, L. Wolde, B. Tadesse, G. Demisie et al. CIMMYT, Addis Ababa, Ethiopia.

Messing, S. A. J., S. B. Gabelli, I. Echeverria, J. T. Vogel, J. C. Guan et al., 2010 Structural insights into maize viviparous14, a key enzyme in the biosynthesis of the phytohormone abscisic acid. Plant Cell 22(9): 2970–2980.

Muzhingi, T., A. S. Langyintuo, L. C. Malaba, and M. Banziger, 2008 Consumer acceptability of yellow maize products in Zimbabwe. Food Policy 33: 352–361.

Neter, J., M. H. Kutner, C. J. Nachtsheim and W. Wasserman, 1996 Applied Linear Statistical Models. McGraw-Hill, Boston.

Owens, B. F., A. E. Lipka, M. Magallanes-Lundback, T. Tiede, C. H. Diepenbrock et al., 2014 A foundation for provitamin A biofortification of maize: genome-wide association and genomic prediction models of carotenoid levels. Genetics 198: 1699–1716.

Pixley, K., N. Palacios Rojas, R. Babu, R. Mutale, R. Surles et al., 2013 Biofortification of maize with provitamin A carotenoids, pp. 271–292, in Carotenoids in Human Health, Nutrition and Health, edited by S. A. Tanumihardjo. Springer Science and Business Media, New York.

Price, A. L., N. J. Patterson, R. M. Plenge, M. E. Weinblatt, N. A. Shadick et al., 2006 Principal components analysis corrects for stratification in genome-wide association studies. Nat. Genet. 38: 904–909.

Qaim, M., A. Stein, and J. V. Meenakshi, 2007 Economics of biofortification. Agr. Econ. 37:119–133.

R Core Team, 2018 R: A language and environment for statistical computing. R Foundation for Statistical Computing, Vienna, Austria.

Robertson, D. S., 1955 The genetics of vivipary in maize. Genetics 40(5): 745–760.

Rodríguez-Concepción, M., and A. Boronat, 2002 Elucidation of the methylerythritol phosphate pathway for isoprenoid biosynthesis in bacteria and plastids. A metabolic milestone achieved through genomics. Plant Physiol. 130: 1079–1089.

Rodríguez-Concepción, M., N. Campos, A. Ferrer, and A. Boronat, 2013 Biosynthesis of isoprenoid precursors in Arabidopsis, pp. 439–456 in Isoprenoid Synthesis in Plants and Microorganisms, edited by T. J. Bach and M. Rohmer. Springer, New York.

Romay, M. C., M. J. Millard, J. C. Glaubitz, J. A. Peiffer, K. L. Swarts et al., 2013 Comprehensive genotyping of the USA national maize inbred seed bank. Genome Biol. 14: R55.

Schwartz, S. H., B. C. Tan, D. A. Gage, J. A. Zeevaart, and D. R. McCarty, 1997 Specific oxidative cleavage of carotenoids by VP14 of maize. Science 276(5320): 1872–1874.

Schwartz, S. H., X. Qin, and J. A. D. Zeevaart, 2001 Characterization of a novel carotenoid cleavage dioxygenase from plants. J. Biol. Chem 276: 25208–25211.

Schwarz, G. 1978. Estimating the dimension of a model. Ann. Stat. 6: 461–464.

Simpungwe, E., T. Dhliwayo, M. Palenberg, V. Taleon, E. Birol et al., 2017 Orange maize in Zambia: crop development and delivery experience. Afr. J. Food Ag. Nutr. Dev. 17(2): 11973–11999.

Singh, M., P. E. Lewis, K. Hardeman, L. Bai, J. K. C. Rose, et al., 2003 Activator mutagenesis of the pink scutellum1/viviparous7 locus of maize. Plant Cell 15(4): 874–884.

Stevens, G. A., J. E. Bennett, Q. Hennocq, Y. Lu, L. M. De-Regil et al., 2015 Trends and mortality effects of vitamin A deficiency in children in 138 low-income and middle-income countries between 1991 and 2013: a pooled analysis of population-based surveys. Lancet Glob. Health 3: e528–e536.

Sun, G., C. Zhu, M. H. Kramer, S. S. Yang, W. Song et al., 2010 Variation explained in mixed-model association mapping. Heredity 105: 333–340.

Sun, Z., J. Hans, M. H. Walter, R. Matusova, J. Beekwilder et al., 2008 Cloning and characterisation of a maize carotenoid cleavage dioxygenase (ZmCCD1) and its involvement in the biosynthesis of apocarotenoids with various roles in mutualistic and parasitic interactions. Planta 228(5): 789–801.

Takahata, Y., T. Noda, and T. Nagata, 1993 HPLC determination of β-carotene content of sweet potato cultivars and its relationship with color values. Japan. J. Breed. 43: 421–427.

Tan, B. C., J. C. Guan, S. Ding, S. Wu, J. W. Saunders et al., 2017 Structure and origin of the White Cap locus and its role in evolution of grain color in maize. Genetics 206(1): 135–150.

Tan, B. C., S. H. Schwartz, J. A. D. Zeevaart, and D. R. McCarty, 1997 Genetic control of abscisic acid biosynthesis in maize. Proc. Natl Acad. Sci. 94(22): 12235–12240.

VanRaden, P. M, 2008 Efficient methods to compute genomic predictions. J. Dairy Sci. 91: 4414–4423.

Vallabhaneni, R., L. M. Bradbury, and E. T. Wurtzel, 2010 The carotenoid dioxygenase gene family in maize, sorghum, and rice. Arch. Biochem. Biophys. 504: 104–111.

Vogel, J. T., B. C. Tan, D. R. McCarty, and H. J. Klee, 2008 The carotenoid cleavage dioxygenase 1 enzyme has broad substrate specificity, cleaving multiple carotenoids at two different bond positions. J. Biol. Chem. 283: 11364–11373.

Vranová, E., D. Coman, and W. Gruissem, 2013 Network analysis of the MVA and MEP pathways for isoprenoid synthesis. Annu. Rev. Plant Biol. 64: 665–700.

Weber, E. J., 1987 Carotenoids and tocols of corn grain determined by HPLC. J. Am. Oil Chem. Soc. 64: 1129–1134.

Welch, R. M., and R. D. Graham, 1999 A new paradigm for world agriculture: meeting human needs. Productive, sustainable, nutritious. Field Crop Res. 60: 1–10.

Yan, J., C. B. Kandianis, C. E. Harjes, L. Bai, E. Kim et al., 2010 Rare genetic variation at Zea mays crtRB1 increases β-carotene in maize grain. Nat. Genet. 42: 322–327.

Zhang, C., X. Chen, R. Zou, K. Zhou, G. Stephanopoulos, et al., 2013 Combining genotype improvement and statistical media optimization for isoprenoid production in E. coli. PLoS ONE 8(10): e75164.

Zhang, Z., E. Ersoz, C. Q. Lai, R. J. Todhunter, H. K. Tiwari et al., 2010 Mixed linear model approach adapted for genome-wide association studies. Nat. Genet. 42: 355–360.

Zhao, Y., H. Sun, Y. Wang, Y. Pu, F. Kong et al., 2013 QTL mapping for the color, carotenoids and polyphenol oxidase activity of flour in recombinant inbred lines of wheat. Aust. J. Crop Sci. 7: 328–337.

Zhu, C., S. Naqvi, J. Breitenbach, G. Sandmann, P. Christou, et al. 2008 Combinatorial genetic transformation generates a library of metabolic phenotypes for the carotenoid pathway in maize. Proc. Natl Acad. Sci. 105(47): 18232–18237.

